# Adaptive scans allow 3D-targeted laser-dissection to probe the mechanics of cell sheets

**DOI:** 10.1101/2022.01.30.478374

**Authors:** Huicheng Meng, Dmitry Nuzhdin, Miguel Sison, Frédéric Galland, Loïc LeGoff

## Abstract

The mechanical actuation of cells by active forces from the cytoskeleton drives tissue morphogenesis. To understand these forces, multicellular laser dissection has become an essential tool for severing tissue locally and inferring tension from the recoil of surrounding structures. However, conventional laser dissection is limited by 2D steering, which is inadequate for embryos and developing tissues that are intrinsically 3D structures. In this study, we introduce a flexible near-infrared (NIR) fs-pulsed laser dissection system that allows for dissection trajectories to proceed in 3D and adapt to the curved surfaces of cell sheets, which are prominent structures in embryos. Trajectories are computed through an unsupervised search for the surface of interest. Using this technique, we demonstrate sectioning of multicellular domains on curved tissue, which was not possible with regular NIR laser scanning.

We apply the developed strategy to map mechanical stresses in the imaginal disc of the developing Drosophila wing. Our targeted, adaptive scans can be used in other non-linear processes, such as two-photon fluorescence imaging or optogenetics. Overall, this new laser dissection system offers an innovative solution for studying complex 3D structures and their mechanical properties.

## INTRODUCTION

Our understanding of morphogenetic processes has considerably improved in the past decade owing to a multidisciplinary effort to address biological shapes in mechanical terms^1–3^. A critical goal is to relate distortions and flows within tissues with patterns of forces imposed by active cytoskeletal elements. From a technical point of view, strains and flows within tissues can be obtained through live imaging of entire embryos at subcellular resolution^4^. Mechanical stresses are more difficult to measure as they require the introduction of a calibrated sensor, in the form of genetically encoded elastic proteins^5, 6^ or calibrated bead sensors^7, 8^. When the introduction of a force transducer within the tissue is not possible, laser dissection (LD) has emerged as a convenient alternative to quantify mechanical stress^9–14^.

The working principle of LD is to locally cut the biological tissues using either UV light or fs-pulsed near-infrared light (NIR). Upon local disruption, the amplitude and speed of the recoil instruct on the mechanical stress that was born by the ablated region^15^. In essence, laser dissections recapitulate at a microscopic scale and without the need for direct physical access the established technique of hole-drilling residual stress measurements, a technique used by Langer, 150 years ago, to investigate the tension of the human skin^16^. The size of the severing depends on the characteristic scale at which tissue mechanics must be probed. Changes in the shape of tissues are well described by continuum modeling approaches at spatial scales larger than the cell^17, 18^. In this context, it is important to probe tissue mechanics at such mesoscopic scales, which is why multi-cellular dissections have been developed using circle or line cuts^19^.

In the field of developmental morphogenesis, multicellular laser cuts could hold great potential, as they allow measurement of the mechanical state of embryonic tissues which are otherwise hard to access from a mechanical point of view. However, the technique is only occasionally used^20, 21^, as it faces one major technical difficulty: most laser steering systems only scan in 2D while embryos and developing tissues are usually curved 3D structures. For example, embryonic tissues are often organized in cell sheets, called epithelia, which take a central role in structuring embryos and precursors of adult tissues^22^. A lot of the forces underlying morphogenesis concentrate at the apical and basal surfaces of these cell sheets, which are sites of enrichment in the force-generating cytoskeleton. The subtle balances or forces along either of these surfaces drive morphogenesis of many developing tissues. It is thus crucial to properly target the dissection laser along these rapidly changing surfaces, which laser-dissection systems could not do so far.

In this methods paper, we present a fs-laser dissection system that dissects epithelial cell sheets in a three-dimensional fashion, effectively adapting to their curved surfaces (Fig. 1). Trajectories are computed through an unsupervised search for the surface of interest. This system allows to perform multicellular cuts on complex curved epithelia, such as *Drosophila* imaginal discs, which are the developing precursors of Drosophila tissues and where conventional 2D systems fail to proceed.

**FIG. 1.**
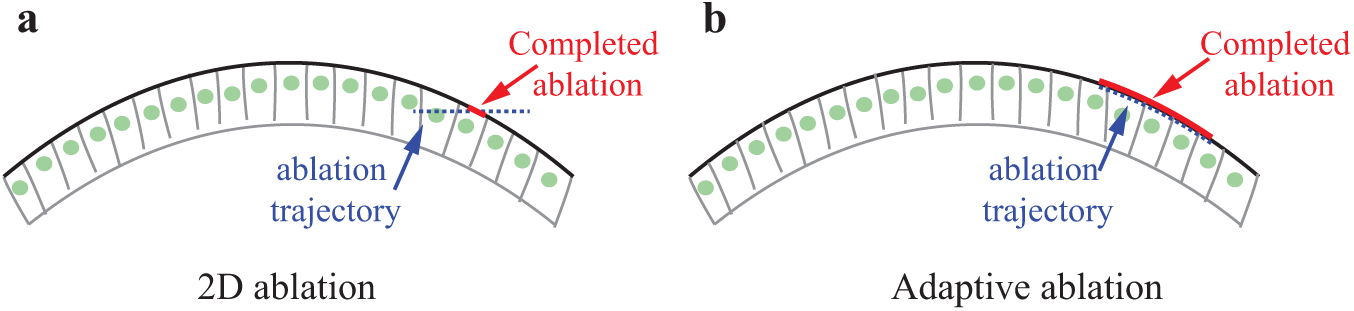
Ablating a curved cell sheet. (a) In a conventional dissection system, the laser focus is scanned along a 2D trajectory which will miss regions of the structure to ablate that are above and below the focal plane of the microscope objective. (b) In an adaptive dissection system, synchronization of the x,y,z steering of the laser ensures that a 3D cut trajectory accurately follows the contours of the structure of interest.

In this study, we employ linear fluorescence imaging using a spinning disc confocal system to visualize tissues. This technique enables rapid imaging with high resolution, making it ideal for capturing epithelial structures located at the surface of embryos, where a shallow imaging depth suffices. By adopting this approach, we align ourselves with the widely accepted configuration for mechanical investigation of embryos, which involves imaging with a spinning disc and dissecting using a fs-pulsed NIR system. Alternatively, some investigators opt to perform laser dissection experiments on stand-alone two-photon microscopes, where both imaging and dissection are carried out using the fs-pulsed NIR laser. However, this setup results in slower and less resolved imaging, which can pose challenges when estimating the fast-changing surfaces of epithelial tissues. We strived to develop and approch that can be adapted to both instrumental contexts.

The structure of the paper is the following : we first present the instrument - which integrates spinning disc imaging with NIR laser dissection in a straightforward manner. We then illustrate the inadequacy of conventional laser steering methods for dissecting embryonic structures, and propose a viable solution by employing targeted adaptive 3D scans along the regions of interest. Finally, we demonstrate our approach by investigating tensions in the Drosophila wing imaginal disc, thereby establishing the efficacy of our methodology.

## RESULTS

### Experimental set-up

The custom-built setup performs imaging and laser dissection in an instrumental configuration which is widely used in developmental morphogenesis. It comprises two optical paths, one for imaging and one for dissection (see methods and Fig. 2a). Briefly, imaging is performed with a spinning disc confocal combined to an inverted microscope using 488 nm and 561 nm lines. A fs-pulsed fiber laser centered at 930 nm is focused in the imaging plane by the objective and steered by galvanometer mirrors placed in a plane conjugated to the back focal plane of the objective. A piezo-actuated stage ensures z-translation of the sample synchronously with the x,y-steering from the scanner to provide 3D trajectories. A data acquisition board outputs properly timed analog signals for x,y,z-translations and their synchronization with the camera and on/off modulation of the lasers. All hardware is controlled using Matlab for maximum flexibility. Specifically, imaging hardware is controlled within Matlab through the *µ*Manager core^23^.

**FIG. 2.**
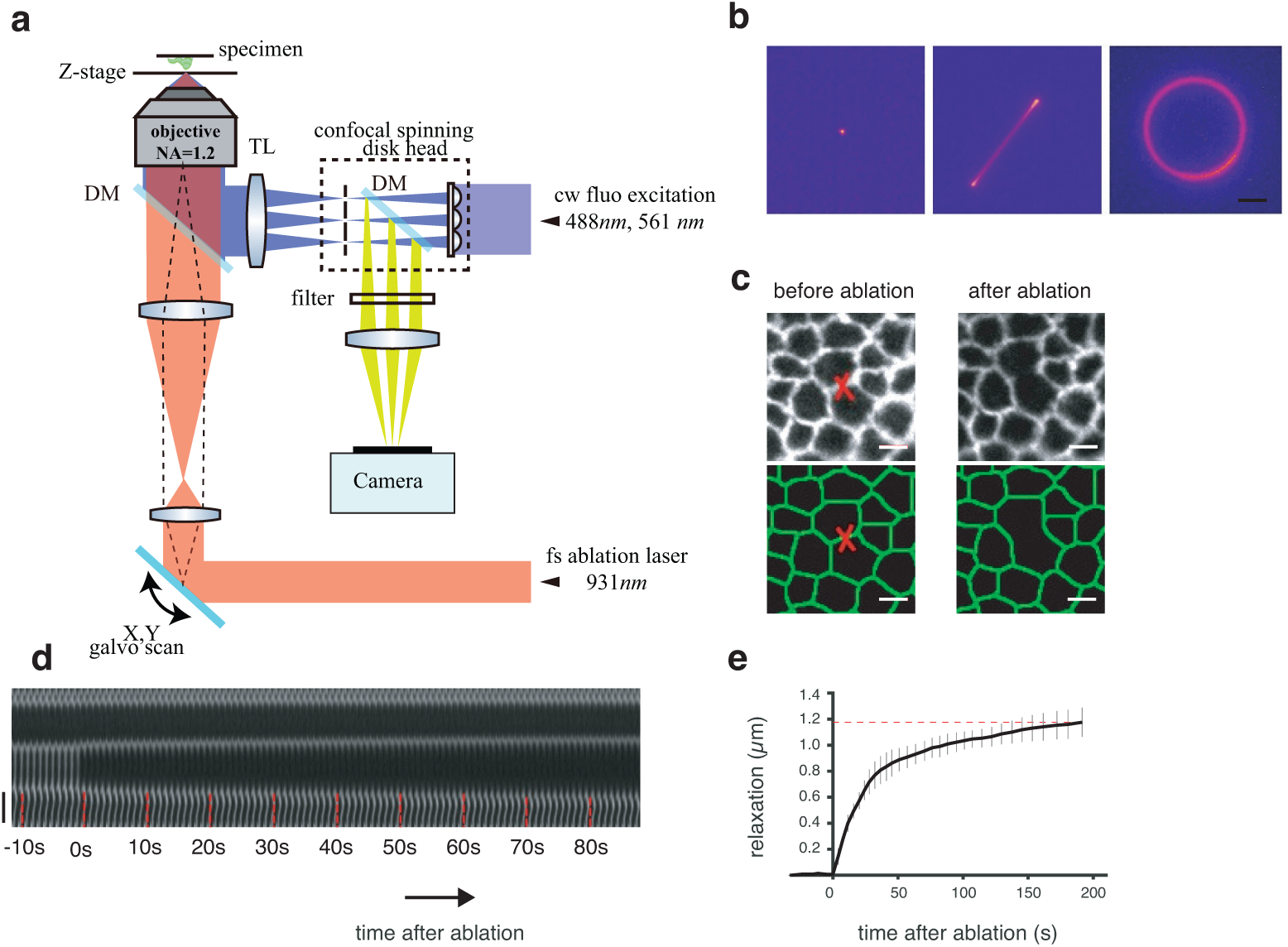
Optical setup for the combined imaging and dissection of epithelial tissues. (a) simplified scheme of the optical layout. DM: dichroic mirror; TL: tube lens (b) Modes of fs-pulsed laser steering for dissection; from left to right: points, line, circles. (c-e) Example of the severing of a single junction. (c) Before/after comparison of the tissue around the severed junction (indicated by a red cross). Cell junctions are visualized through the imaging of a cadherin:GFP fusion. The bottom panel shows the segmented cell outlines. (d) Relaxation dynamics of the severed junction as observed through a kymograph. (e) Quantification of the relaxation. All scale bars are 3*µm*

The resulting apparatus performs 2D scanning severing along points, lines or circles (Fig. 2b) in a similar fashion to previously published works^9, 10, 12, 17, 19, 24^. The focused laser delivers a pulse peak power density near 7.42×10^12^W*·cm^−^*^2^. With a pulse duration of 130 fs, the system has the prerequisite to perform plasma-induced ablation^15, 25^. A typical experiment is illustrated in Fig. 2c-e. Individual cell-cell epithelial junctions are severed through a point or a small line severing (Fig. 2c) leading to the mechanical relaxation of the surrounding tissue (Fig. 2d,e). A detailed analysis of the relaxation curve after severing requires taking into account the material properties of the tissue and the boundary conditions of the severed domain^17, 20^. In the present study, we simply focus on measuring the tension born by the tissue before the cut. In this context, a simple Kelvin-Voigt model reveals that the initial slope of the relaxation curve instructs on the tension to friction ratio *T*_0_*/ζ*, where *T*_0_ is the tension initially born by the junction and *ζ* is the effective friction of the medium^26^ (see also the analysis provided in Appendix A).

### Failed multi-cellular dissections with 2D scans

Developing tissues are often curved structures. Such is the case of the *Drosophila* wing imaginal disc used in this study^27, 28^ -an important model system to study the regulation of growth and morphogenesis^29^. In this context, performing cuts at intermediate to large scales is difficult as 2D laser steering cannot adapt to the 3D contour of the surface. We could perform multi-cellular cuts at the apex of the tissue - provided the tissue is sufficiently flat and horizontal there (Fig. 3a-d). The trajectory of the cut then follows the contours of the tissue (red circle in Fig. 3a). For demonstrative purpose, we assess the trajectory by scanning the NIR laser at power < 50mW. At such low power, no ablation is generated, but the marker (Ecad:GFP) emits two-photon fluorescence that we can detect. We can then ensure that the laser is properly focused on the targeted surface throughout the 3D scan. Indeed, fluorescence is homogeneous along the circular trajectory (Fig. 3b). A complete circular cut proceeds if we scan the laser at high power (*∼*165 mW) along the same trajectory (Fig. 3c,d). Successful cuts, however, are limited to a small apical region of the tissue. In slopped regions, the 2D trajectory cannot follow the contours of the tissue (red circle in Fig. 3e). At low NIR power, the emitted fluorescence of the marker is non-homogeneous (arrows in Fig. 3f). Ablation, which relies on a multi-photon ionization process^25^, is very sensitive to the laser focus being localized at the adherens plane. We observe only partial cuts (arrows in Fig. 3g,h) when scanning a high-power laser along the 2D circular trajectory as ablation is effective in only a fraction of the trajectory (arrows in Fig. 3h).

**FIG. 3.**
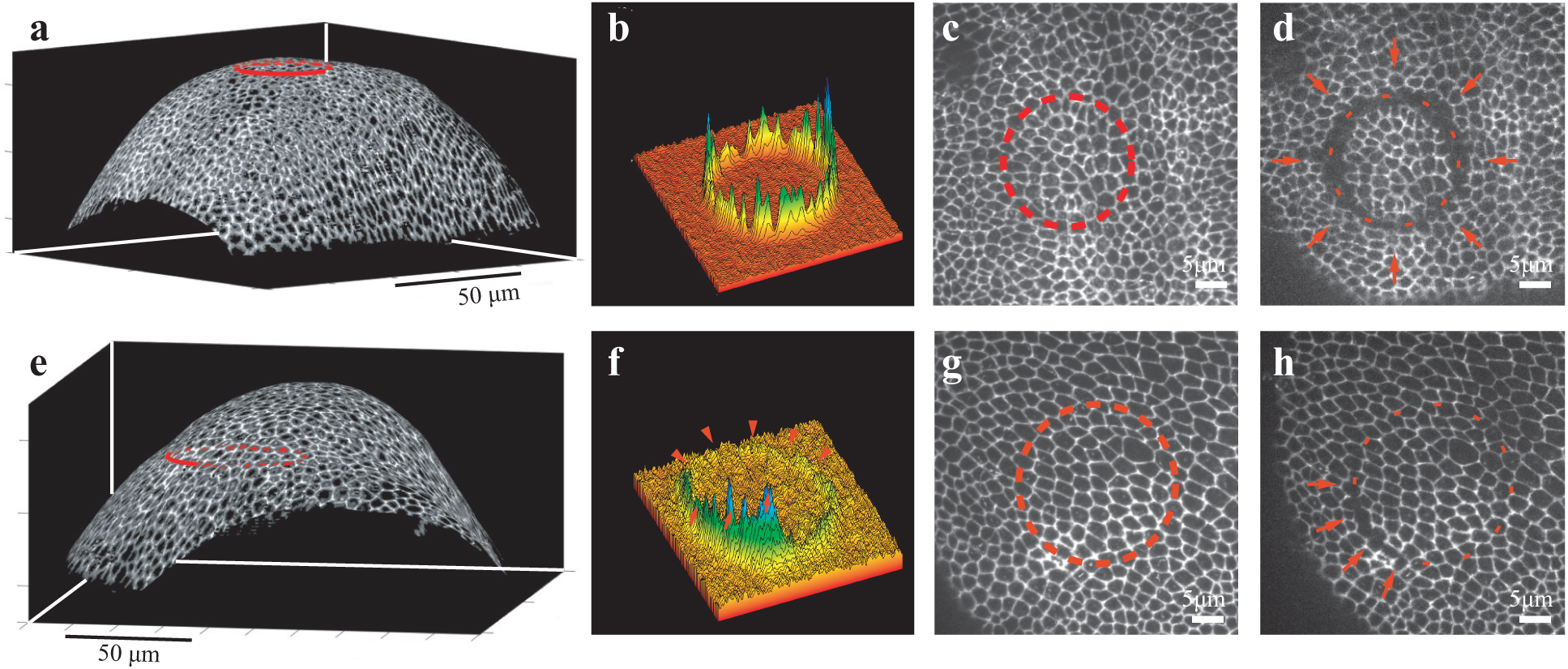
Failed circular cuts on curved tissues. (a-d): 2D cuts along the flat apex of the tissue are successful. (a): 3D topography of a curved wing imaginal disc (cadherin:GFP labeling). A 2D circular trajectory (red line) can follow the contours of the tissue. (b): At low excitation power, the laser can excite GFP-fluorescence of the junctional marker on the whole trajectory through a two-photon excitation effect. (c,d): Using the same laser along the same trajectory, but at high power, severs the tissue on the whole trajectory (red arrows). (e-h): 2D severing along the sloped regions of the tissue fails. (e): The 2D circular trajectory does not follow the contour of the surface. (f): At low excitation power, the laser excites fluorescence of the junctional marker only partially, failing to do so in large portions of the trajectory (red arrowheads). (g,h): The laser at high power along the same trajectory severs the tissue in a fraction of the trajectory (red arrows), failing to do so elsewhere.

### 3D dissections with adaptive scans

We implemented a 3D dissection system to overcome the observed failure of multicellular severing with a planar trajectory (Fig. 4). The principle is first to estimate the surface of the epithelium *Z_s_*(*x, y*) through image analysis (see details below and in the methods). We then generate a trajectory (*x*(*t*)*, y*(*t*)*, z*(*t*)), enforcing that it lies on the computed surface *Z_s_*(*x, y*). In the course of dissection, the z-component of the trajectory is generated by moving the sample stage in synchrony with the xy-actuation from the galvanometric mirrors (Fig. 4a,b). The proposed 3D dissection rely on an estimation of the epithelial surface *Z_s_*(*x, y*), which we briefly present here (see details in the methods section). The surface that we target for severing corresponds to the adherens plane of the epithelial cells which concentrate a lot of cytoskeletal activity underlying morphogenesis^22^. We identify the adherens plane through its enrichment in the protein fusion E-cadherin:GFP. In the first step, we detect bright voxels in the volume, *ie* voxels that might contain GFP signal, assuming a normalized intensity distribution for the non-fluorescent voxels (background) after correction of signal inhomogeneities. The estimation then relies on piecewise polynomial fits of the surface. These fits rely on a RANSAC approach^30^, which classifies the detected bright voxels into a set of inliers (bright voxels estimated to be part of the surface) and outliers (bright voxels estimated not to be part of the surface). This procedure maximizes the number of inliers, which amounts to finding the polynomial surface which is the most densely populated in bright voxels. It ensures the robustness of the estimation to noise and to the presence of many fluorescent voxels located outside of the main surface (outliers). When we estimate only a small portion of the surface (typically the size of a multi-cellular annulus *∼* 15*µm*), a single polynomial is enough to locally fit the surface (Fig. 4c). When we estimate a large surface (typically the entire epithelial surface), the RANSAC polynomial fits are performed on overlapping windows, and the different reconstructions are fused (Fig. 4d, see methods).

**FIG. 4.**
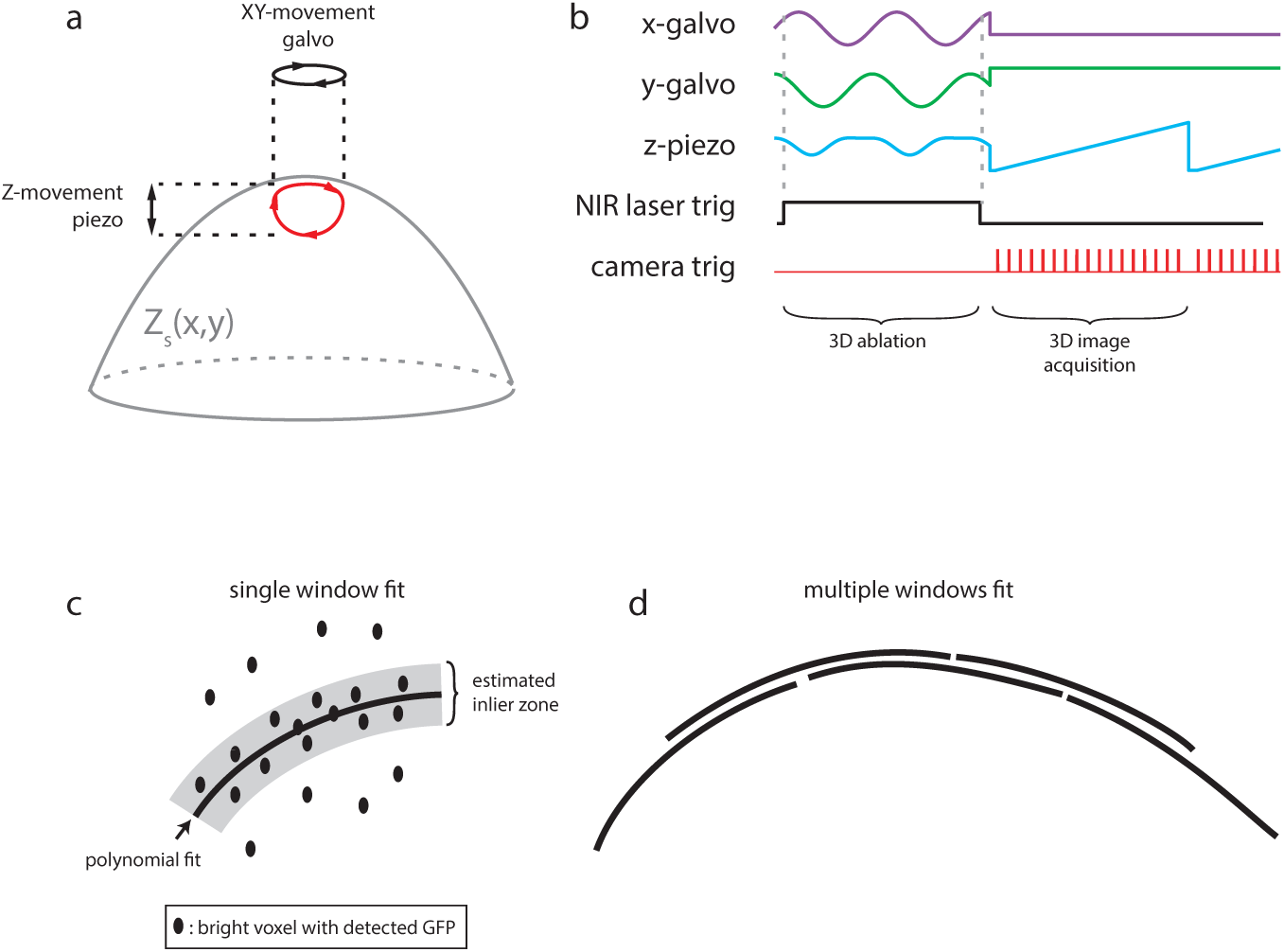
Principle of 3D dissections. (a) A trajectory (*x*(*t*)*, y*(*t*)*, z*(*t*)) is computed so that it lies on the estimated surface *Z_s_*(*x, y*). XY-translation is performed with galvanometric scanners, while Z-translation is performed with the translation stage of the microscope, or alternatively a focus control such as a deformable mirror or an electrically tunable lens. (b) Experimentally, the 3D trajectory relies on the synchronous actuation of *x, y, z* translation as well as shutter of the NIR laser. The dissection phase is followed by a fast 3D imaging phase to measure the displacements induced by the cut. (c,d) Estimation of the surface of interest *Z_s_*(*x, y*). (c) On short-length scales, the surface is fitted with a second-order polynomial. With the RANSAC approach, the fit maximizes the number of bright voxels in its vicinity (estimated inlier zone on the scheme) (c) On larger length scales, the surface is fitted with multiple overlapping windows, each window using the RANSAC approach of (c).

With such a process, we could perform multi-cellular cuts in sloped regions of the wing imaginal disc (Fig. 5a,b). When scanning the NIR laser at low power (no ablation) in a circular trajectory on the surface, the generated fluorescence is homogeneous along the trajectory (Fig. 5c). At high laser power, the induced cut is also performed homogeneously along the trajectory (Fig. 5b,d). When performing circular cuts, the trajectories are made circular locally along the tangent plane of the surface. To do so, we delineate the circle of given radius on the local tangent and project this circle on the surface along the normal direction to ensure maximal confocality of the dissection trajectory and the surface (see methods and Fig. S4).

**FIG. 5.**
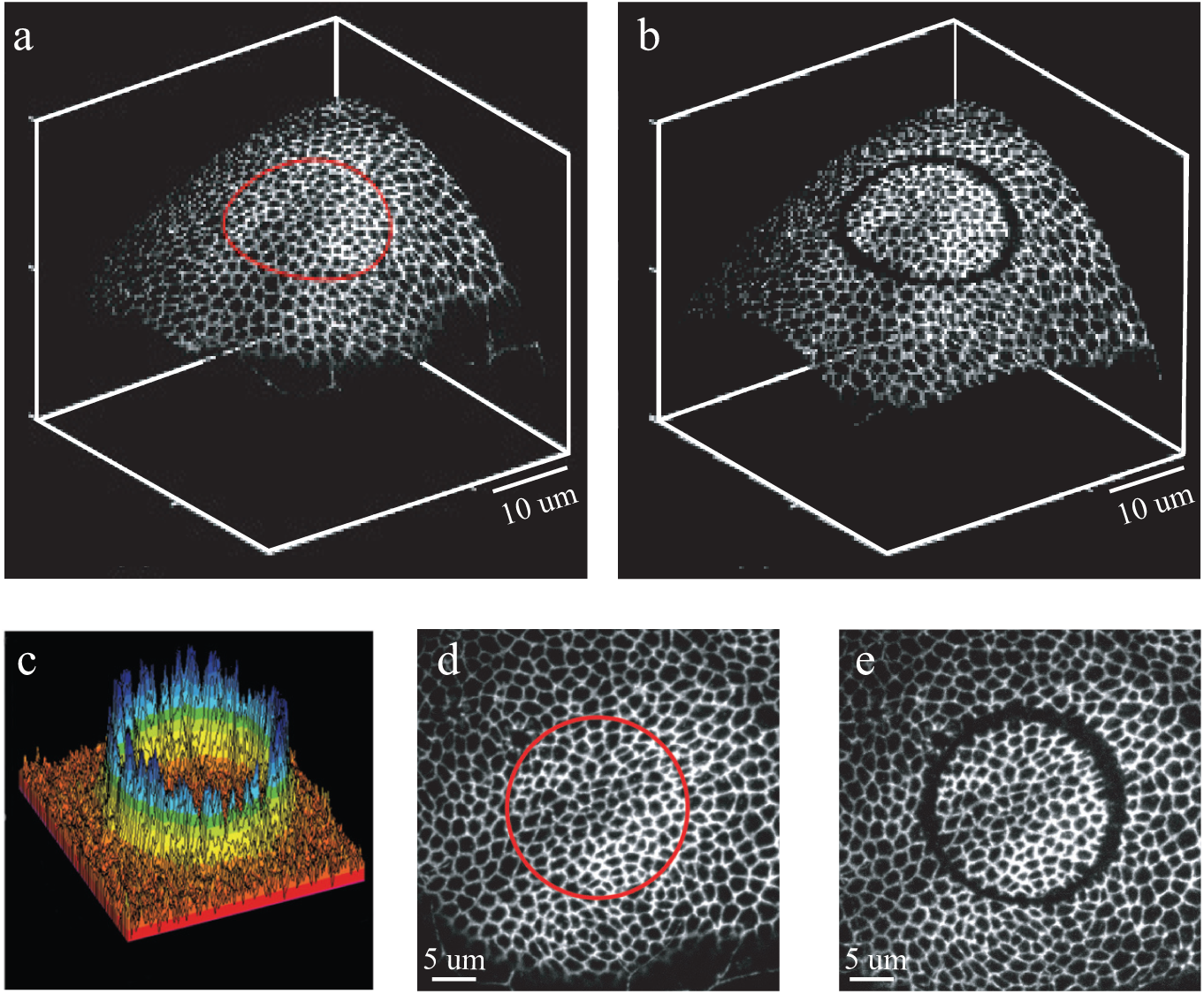
Example of 3D dissections. (a,b) 3D representation of a wing imaginal disc (close-up view) before(a) and after (b) a multicellular cut in a sloped region. (c) At low excitation power, two-photon fluorescence is emitted homogeneously along the 3D trajectory. (d,e) 2D visualization of the dissection process shows that severing was effective along the entire trajectory.

In our ablation setup, the surface is estimated from the data-rich 3D stacks acquired using a spinning disc confocal microscope. As discussed in the introduction, spinning disc confocals are widely used for imaging embryonic epithelia. Alternatively, investigators may use stand alone two-photon microsopes, which use the fs-NIR laser both for imaging and dissection -at different laser power. Can our adaptive 3D dissection approach be adapted to such a standalone fs-NIR laser system? The slower imaging speed associated with two-photon fluorescence microscopes might hinder the effectiveness of the adaptive strategy. Additionally, lower photon fluxes and different signal statistics could affect the accuracy of surface estimation. We demonstrate the adaptability of our approach in Appendix B, where we propose an accelerated method for rapid surface estimation with a single-point two-photon microscope. This method involves performing a highly expedited fractional scan of the volume using Lissajous trajectories, resulting in a ten-fold acceleration compared to a raster scan. The same surface fit, utilizing a robust approach, can then be employed to work with such sparse and noisy data.

### Application: mapping mechanical asymmetries in the wing imaginal disc

We use the ablation setup described in Fig 2a to explore the distribution of tensions in the wing imaginal discs. The tissue is under patterned mechanical stress, which influences growth and the organization of the cytoskeleton within cells^31, 32^. Tension in the wing imaginal disc has mostly been investigated via single junction, point ablations. Multi-cellular ring dissections offer an attractive alternative as they assess a mesoscopic scale that translates naturally to continuum mechanics and provides robust measurements by averaging the response of multiple cells^17, 19, 21^.

Most importantly, circular dissections provide a symmetric perturbation that allows revealing most clearly the asymmetries inherent to the probed tissue: the main axis of relaxation after severing provides a measurement of the main axis of tension of the tissue, locally. Figure 6a-c shows the tissue before and after the circular cut is performed. The group of cells isolated by the cut contracts under the influence of their constitutive tension (inner wound margin in yellow in Fig. 6b,c). The outer wound margin (blue in Fig. 6b,c) expands orthogonally due to the constitutive tension of the surrounding tissue. The inner and outer margins have their major axis orthogonal to each other. Figure 6d maps the displacements of the surrounding tissue following dissection, while Fig. 6e,f quantifies the opening of the wound along the major and minor axis of the outer wound. A detailed analysis of the relaxations after similar annular severings was performed by Bonnet *et al.*^19^. We resort to a simpler analysis focusing only on the asymmetries in tensions: we use the ratio of the slope at the origin of the relaxations as a proxy for the ratio of the tensions.

**FIG. 6.**
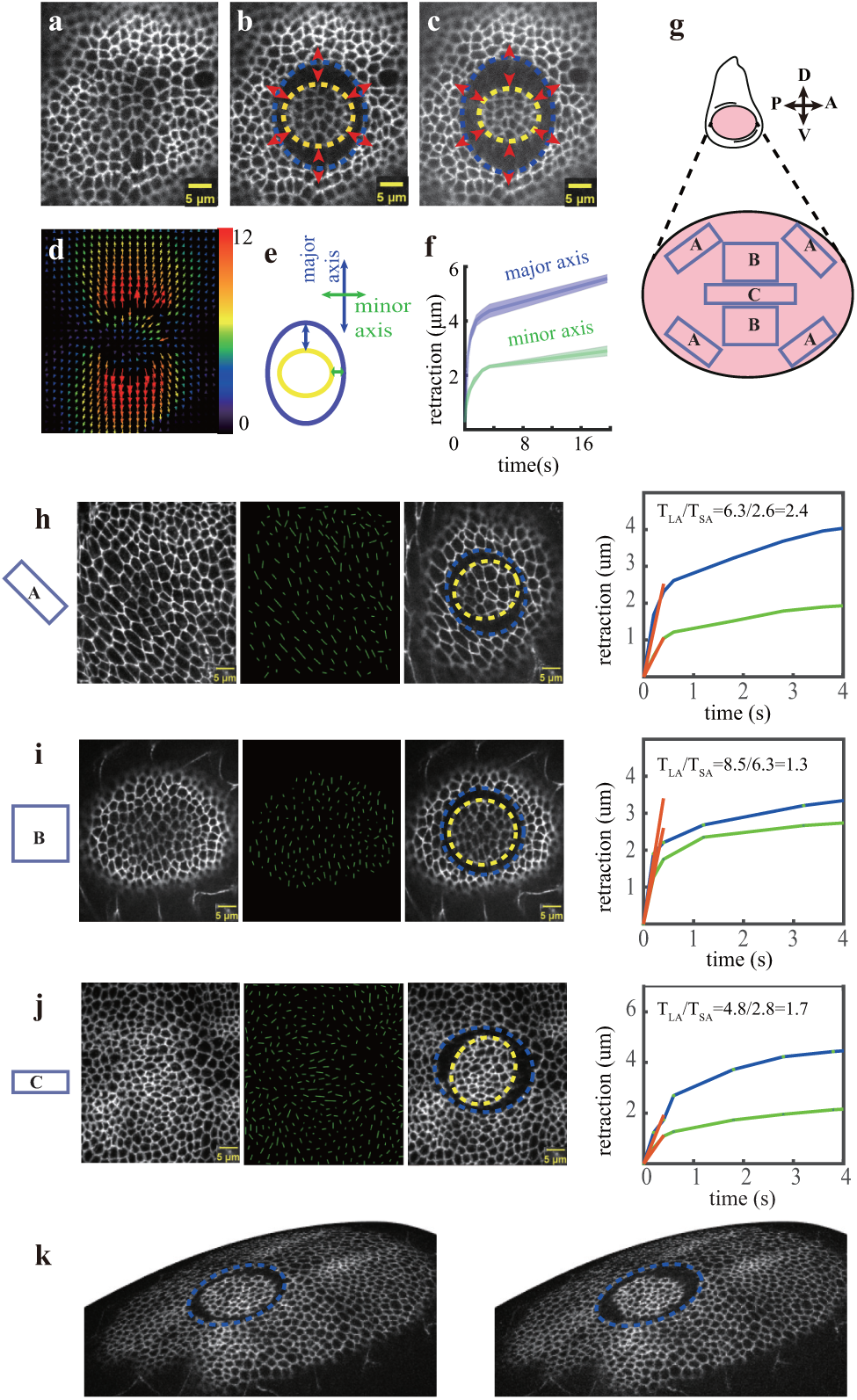
Mapping mechanical asymmetries in the wing imaginal disc. (a-f) example of a dissection showing the tissue before (a) and after (b,c) a circular dissection. (d) Displacement map after dissection using particle image velocimetry. (e,f) quantification of the opening of the wound along the outer wound boundary long and short axis: scheme in (e) and measurements in (f). (g) Regionalization of the wing imaginal disc into peripheral (region A), medial (region B), and margin domains (region C). In the scheme of the wing imaginal disc on top, the presumptive wing is colored pink while other tissues are in white. (h-j) Examples of dissections in the mentioned domains. (h) In the periphery (region A), cells are anisotropic and oriented along the tangent of the tissue (see green segments indicating the major axis of cells). The main axis of the outer wound is oriented along that same direction, and the response is markedly anisotropic with a ratio *T_LA_/T_SA_ ∼* 2.4. (i) In the medial regions (region B), cells are more isotropic and a mean orientation does not appear clearly. The relaxations are also more isotropic with a ratio *T_LA_/T_SA_ ∼* 1.3. (j) In the margin (region C), despite the fact that cells are not particularly anisotropic, the response after dissections is anisotropic (*T_LA_/T_SA_ ∼* 1.7) with the main axis oriented along the margin. (k) 3D representation of the same tissue as (j), at two-time points following the1d1issection.

We performed dissections in different regions of the wing imaginal disc. Following LeGoff *et al.*^31^, we distinguish peripheral regions (region A in Fig. 6g), medial regions (region B in Fig. 6g), and the margin (region C in Fig. 6g). The peripheral and margin regions experience anisotropic stress, while the medial regions are isotropic. We observed characteristic behavior in different regions of the wing disc (Fig. 6h-j). Figure 6h shows the response of the tissue in the periphery. As described in^31, 32^, cells are anisotropic in this region (see the major axis of cells represented by green lines) and globally oriented along the tangent of the tissue. We observe an anisotropic relaxation of the wound, with the main axis of stress qualitatively oriented along the same direction as the cells. To characterize the anisotropy, we measure the slope ratio at origin along the major and minor axis *T_LA_/T_SA_ ∼* 2.4. In the medial region, the cells are much less anisotropic and a global orientation is hardly visible. The tissue there relaxes in a fairly symmetric fashion *T_LA_/T_SA_ ∼* 1.3 (Fig. 6i). In the margin domain (region C), the relaxation is also anisotropic, with the major axis oriented along the margin, albeit to a less extent than in the periphery *T_LA_/T_SA_ ∼* 1.7. The anisotropic relaxations at the periphery and at the margin betray an anisotropic stress which is in agreement with previous work^21, 31, 32^. Lastly, we stress the importance of the 3D trajectory for the ring dissections. As can be seen in the 3D rendering of Fig. 6k, the 3D adaptive scan is necessary to ablate along the sloped topography of the tissue (see also supplementary movie 1).

## DISCUSSION

We have designed a system to perform precise laser cuts on curved biological surfaces using adaptive trajectories along the biological surface of interest. The surface of interest is first estimated using high content 3D-information from a spinning disc confocal. Then a fs-pulsed laser is scanned along a 3D trajectory on the surface to generate a local ablation that severs the tissue.

Laser dissections have become a central tool in mechano-biology to infer tensions in tissues from the recoil velocity of the severed structures^9–15, 33, 34^. In particular, multi-cellular cuts probe mesoscopic scales which are well suited for the confrontation with continuum descriptions, the natural language of mechanics. Circular dissections are particularly convenient as their symmetry directly reveals the anisotropy of tension from the shape of the relaxation. Among the various strategies to generate the cuts, a method of choice is to use NIR, fs-pulsed lasers to induce a plasma which is well controlled in space. To mitigate the difficulty of performing multicellular cuts on curved tissues, we have introduced an adaptive strategy where the surface of interest is first estimated around the region of interest, and the laser is then scanned along a computed 3D trajectory on the surface. Multi-cellular dissections have already been performed on the wing imaginal disc or other curved tissues^21^. However, it requires a lot of trials and errors to position the region of interest flat with respect to the focal plane of the imaging objective. We hope our approach will help generalize this powerful approach to probe stresses within biological tissues. We designed our analysis pipeline so that it could be easily adapted to other imaging contexts, as demonstrated in Appendix B and Fig. S1. Indeed, many NIR fs-pulsed lasers are only available in stand-alone non-linear imaging setups. There, the single-point scan and the low photon flux of non-linear contrasts make the 3D scan of the complete tissue problematically slow, as the tissue may move by the time the volume is scanned and the surface estimated. We thus adapted the experimental approach using Lissajous scans of the NIR laser to rapidly probe the global features of the tissue and estimate the surface from this fractional scan. Lissajous trajectories are well adapted to scan fractionally at high speed as they don’t introduce unnecessary high harmonics like raster scans do^35, 36^.

While we recently introduced the concept of smart-scans on surfaces for linear fluorescence^28, 37^, we demonstrate here the concept for non-linear processes for the first time. From an instrumental point of view, the targeted illumination of Abouakil *et al.* relied on a spatial light modulator in a wide-field optical configuration. Here, the 3D severing relies on the control of the scan path of a single-point, galvo-based scanning microscope. In the current configuration, our set-up performs 3D scans by synchronizing a rather slow z-stage with the fast xy-galvanometric scans. Faster z-scans could be achieved through the use of an electrically tunable lens^38^ or a deformable mirror^39^.

In this work, we investigate the tensile state of the *Drosophila* wing imaginal disc. Our experimental paradigm will be useful in a broad range of other biological contexts: i-other embryonic structures which are most of the time curved - *eg* the mouse, C. elegans, avian, *Xenopus* embryos; ii-organoids, which are also curved; iii-individual cells, when investigating mechanics of the thin cytoskeletal layer underneath the cell membrane that controls cell shape - the cortex^40^. Our mechanical characterization focuses only on the early response of the tissue upon laser severing to measure the tension to friction ratio, *T*_0_*/ζ* . This is the simple measurement that many investigators focus on^12, 21, 26, 31–33^, but more advanced rheological measurements could also be performed with our technique^17, 19, 41^.

Beyond dissection experiments, the proposed adaptive scans could be used for optogenetic control of cell behavior. Cell contractility controlled by light is a powerful means to investigate tissue mechanics^42^. The optogenetic activation must ideally be targeted to a well-defined spatial domain of cells -*eg* the apical or basal layer. Current approaches use complex genetic schemes to anchor the light-sensitive moiety to specific sub-regions of the cell^43^. Alternatively, our adaptive scans could be used to confine optogenetic activation through a two-photon effect along the surface of interest. In a more applied context, ultrashort pulses are used in surgery and dentistry^44, 45^. Adaptive scans could significantly improve the precision of dissection in these contexts. Lastly, the proposed adaptive scans could be used in a non-linear imaging context, where restriction of the scan path to a thin shell around the surface of interest could be an efficient means to reduce the light dose and speed up imaging, providing a generalization of the geometry-driven scans of Olivier *et al.*^46^.

## MATERIAL AND METHODS

### Spinning disc imaging

The laser dissection set-up combines a spinning disc fluorescence microscope with a fs-pulsed NIR laser for severing (Fig. 2a). The imaging branch of the setup consists of an inverted Nikon Eclipse Ti microscope (Nikon Instruments) equipped with a high NA water immersion objective (Plan Apo 60*X*, NA=1.2, Nikon) and a Yokogawa spinning disk unit (CSU-X1, Yokogawa Electric), with 50*µm* pinhole size. Two excitation lasers (488/561 nm, 20mW, coherent OBIS LX) are combined with a dichroic mirror (561 nm laser BrightLine® single-edge laser dichroic beamsplitter, Semrock) and directed towards the Yokogawa unit after a 10*X* expansion. Fluorescence emitted by the sample and spatially filtered to retrieve sectioning by the spinning disc unit is imaged onto a sCMOS camera (pco.edge, PCO AG, Germany) mounted on the microscope’s left-side port, resulting in a pixel size of 98 nm. Z-scanning is achieved by moving the Z-Pizeo stage (Nano-Z stage, from Mad City Labs). Imaging is controlled in Matlab via a *µ*Manager^23^/Matlab interface. Synchronization with laser dissection is performed with an input/output data acquisition card (USB-6251, National Instruments).

### Laser dissection experiment

Laser-dissection experiments are performed with a near-infrared pulsed (*∼* 130 fs pulses) fiber laser (YLMO 930±10 nm, MenloSystems) operating at 50 MHz. The laser is first expanded, then reflected by a galvanometer-based laser scanning device (6215 H, Cambridge Technology Enterprises) for steering in the sample. The mid-point of the galvanometer mirrors is then relayed onto the back focal plane of the objective used for spinning disc imaging (Plan Apo 60*X*, NA=1.2, Nikon) with a pair of achromatic doublets which also provide a 4*X* magnification with the focal length *f*_1_=75 mm and *f*_2_=300 mm. The laser is then focused by the microscope objective, delivering pulse peak power density near 7.42×10^12^W*·cm^−^*^2^. Two-dimensional laser dissection is obtained by scanning the laser beam over the focal plane with the help of the galvo-scanner. Three-dimensional dissection is achieved moving the Z-Piezo stage (Nano-Z stage, Mad City Labs) and the galvano scanner along the X,Y and Z directions simultaneously (Fig. 4a,b). A data acquisition card (USB-6251, National Instruments) along with Matlab software and *µ*Manager is used for data acquisition and to control the scan mirrors, shutter and Z-stage, all devices are synchronized by the external trigger that we calculated according to the ablative region.

### Software and instrument synchronization

To enable interactive manipulations of the scan path, a custom-written Graphical user interface (MATLAB) has been developed, which allows the user to perform different ablative patterns (point, linear, and 3D circular) directly on the target sample. To select the location of dissection, the total surface is estimated using the procedure described below. The user can then interactively manipulate a 3D rendering of the tissue, generated by projecting signal of the adherens plane onto the 3D surface, to define the dissection pattern. Upon selection of the dissection spot/region, the program re-evaluates the surface of interest in a small window around the region of dissection (see below). It then computes the exact 3D trajectory of the focal spot so that it is as close as possible to the desired trajectory (line, circle) while being confined to the curved surface (see below). Synchronized signals are then generated and sent via the DAQ board to control the galvanometric mirrors, the Z piezoelectric stage, laser, and camera triggers.

### Identification of a surface of interest

We adapt the surface estimation strategy of Abouakil *et al.*^28^. The surface of interest is estimated from the sampling of only a small fraction of the voxels-space, which we call Ω_0_. When using high-content 3D-data from the spinning disc, Ω_0_ is determined by randomly picking voxels in the full volume of acquisition (typically 0.1% of the voxels). We first correct spatial inhomogeneities of the background signal by introducing a normalized signal,

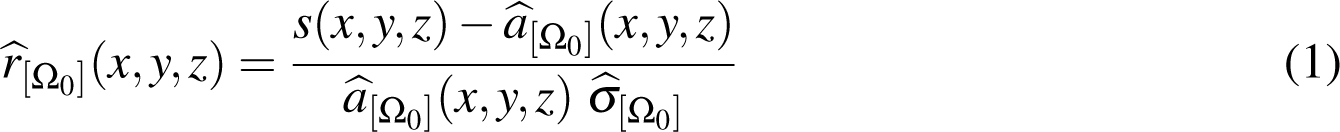

where *s*(*x, y, z*) is the measured signal, *â*_[Ω0]_(*x, y, z*) is an estimation of spatial inhomogeneities of the background and *σ̂*_[Ω0]_ is an estimation of the standard deviation of *s*(*x, y, z*)*/â*_[Ω0]_(*x, y, z*) on the background. Both *â*_[Ω0]_(*x, y, z*) and *σ̂*_[Ω0]_ are determined using only the voxels in Ω_0_. Through this normalization, the histogram of *r̂*_[Ω0]_ falls down on a heavy-tailed Gaussian-like distribution in the case of the spinning disc data (Fig. S3). Assuming the gaussian curve corresponds to the background and the heavy tail to the bright points, we set a threshold *T* from a chosen probability of false alarm (pfa), such that pfa 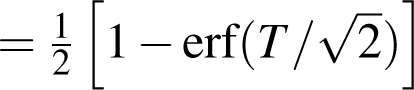, where erf is the error function. We chose a pfa of 1%, which corresponds to *T ≈* 2.33.

In a second step, we interpolate the epithelial surface modeled as *z* = *Z_s_*(*x, y*), using the detected bright points. To cope with noise on the location of bright points and outliers, we use local second-order polynomial fits of the bright point z-coordinates, combined with RANSAC-based outlier removal^30^. When surface estimation is performed in a small region of interest, around the ablated region for example, only one window is used (Fig. S1a), leading to one polynomial fit of the form *z* = *Z_s_*(*x, y*) where:

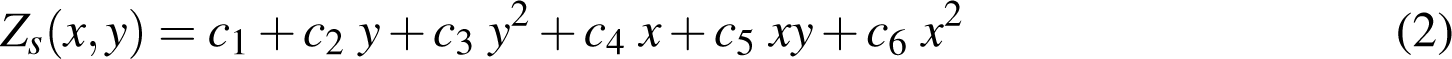

with *c*_1_*, …, c*_6_ the 6 parameters of the surface that have been estimated.

When estimating surface over large regions of interest, for example and entire field of view (*∼* 100*µm*), the fits are estimated in overlapping windows (Fig. 4d) and are then fused. To fuse the estimations in the overlapping regions, we do not use the parameters of the quadratic fits (eq. 2), but instead use the corresponding inliers from the RANSAC approach. More specifically, bright points that are inliers in all of the overlapping quadratic fits are preserved, while those that are inliers in only a fraction (1*/*4, 2*/*4 or 3*/*4) of the overlapping windows are discarded. The surface is then estimated through the interpolation of the remaining inliers using a simple bi-cubic harmonic spline interpolation^47^. Figure S2 shows the successive steps of surface estimation for the tissue of Fig. S1. Once the surface of interest *Ẑ*(*x, y*) is determined, it can be converted into a thin shell by setting a thickness along *z* of *ε* = 3*µm*, which is slightly larger than the thickness of adherens junctions.

### Preparation of biological samples

All observations were performed on living Drosophila tissues, using an E-cadherin:GFP knockin to image adherens junctions^48^. We imaged wing imaginal discs as in LeGoff et al.^31^; Living tissues were dissected from late third instar larva using a stereo-microscope, and cultured in a drop of Grace’s insect medium (sigma) in a glass bottom petri-dish. All experiments are performed at 22*^◦^*C.

## ACKNOWLEDGEMENTS

We thank Sophie Brasselet for discussions on the project, Dana Bruner and Morgane Chauvet for technical help.

This work was funded by the following agencies: Agence Nationale de la Recherche (ANR-18-CE13-028, ANR-17-CE30-0007, ANR-22-CE42-0010) ; Excellence Initiative of Aix-Marseille University - A*Midex (capostromex), a French Investissements d’Avenir programme; This project is funded by the « France 2030 » investment plan managed by the French National Research Agency (ANR-16-CONV-0001, ANR-21-ESRE-0002), and from Excellence Initiative of Aix-Marseille University - A*MIDEX. HM thanks the support of the China Scholarship Council(CSC).

## AUTHOR CONTRIBUTIONS STATEMENT

LLG conceived the experiments. HM built the apparatus with the help of MS, HM conducted most experiments except for two photon microscopy which was conducted by HM and DN. HM, FG, and LLG developed the algorithms and performed data analysis. LLG wrote the paper. All authors reviewed the manuscript.

## DATA AVAILABILITY STATEMENT

Data sets generated during the current study are available from the corresponding author on reasonable request. Matlab implementation of the surface estimation algorithm is available at https://www.fresnel. fr/perso/galland/LSA2021/.

## SUPPLEMENTARY MATERIAL

Appendix A: modeling mechanical relaxation with a simple Kelvin-Voigt model

Appendix B: Rapid surface estimation with a point-scanning microscope using Lissajous scans

Figure S1: Rapid surface estimation in single-point scanning.

Figure S2: Steps of surface estimation

Figure S3: Intensity distributions after correction of inhomogeneities

Figure S4: Geometry of dissection on a curved surface

## Appendix A: modeling mechanical relaxation with a simple Kelvin-Voigt model

To analyse the mechanical relaxation after the laser cut, we use the simple Kelvin-Voigt viscoelastic model, where a spring and a dashpot are arranged in parallel. Such as simple configuration is commonly used to investigate the mechanical response of epithelial tissues^19, 24, 49, 50^ and cell junctions^12, 33, 51, 52^ after laser severing. Assuming a linear slab of tissue tensed by a constant tensile force *T*_0_, we write the mechanical equilibrium for the quasi-static movements:

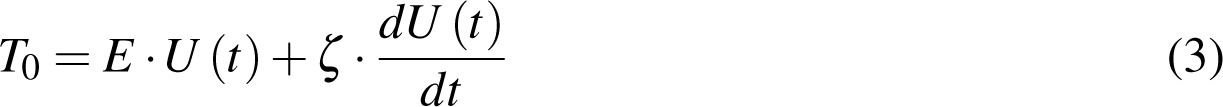

where *t* indicates time, *U* (*t*) is the deformation, E is the spring constant, *ζ* is the viscosity coefficient introduced by the dashpot. Solving for *U* (*t*) (with *U* (*t* = 0) = 0) we find:

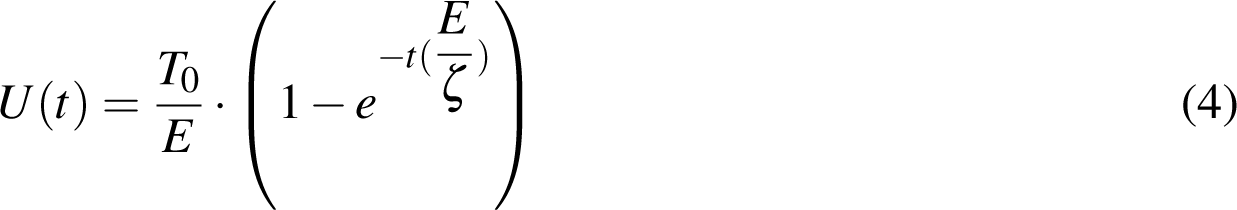

which approximately describes the creep behavior of tissues after dissection. The asymptotic value of *U* (*t*) after laser dissection is proportional to the tensile force initially sustained by the tissue or cell junction with a proportionality set by the Youngs modulus of the tissue (*U_max_* = *T*_0_/*E*). The initial slope of *U* (*t*) after laser dissection is also directly proportional to the tensile force initially sustained by the tissue or cell junction with a proportionality set by the friction (*dU/dt|_t_*_=0_ = *T*_0_*/ζ*).

With fs-pulse NIR laser, the cuts are highly localized. The tissue then suffers minimal damage and recovers very fast after the dissection. Thus the long time response (*t >* 40*s*) tends to be dominated by the rapid sealing of the cytoskeleton, not by the passive viscoelastic response of the tissue. We then prefer to use the initial slope of the relaxation curve to estimate tension.

## Appendix B: Rapid surface estimation with a point-scanning microscope using Lissajous scans

We address the portability of our approach to other instrumental contexts. In our dissection setup, the surface is estimated from the data-rich 3D-stacks of a spinning disc confocal microscope. Most dissection setups, however, exist in the context of stand-alone two photon fluorescence microscopes -not associated with a spinning disc system. A weakness of our approach could then be a poor portability to this usual instrumental context, where the single point scanning and lower photon fluxes make imaging in 3D slower and reduces signal to noise ratio. A fast surface estimation is a requirement, as biological tissues are most of the time dynamic structures. To make our method as general as possible, we specifically designed our surface estimation (see main text and methods) to also work with sparse and noisy data. We then propose an accelerated surface estimation for single point scanning, which consists in performing a very rapid fractional scan of the volume with Lissajous scans. Our surface estimation can handle such a small collection of data points. We experimentally demonstrate the surface estimation with Lissajous scans on a separate, stand-alone two photon setup. For that purpose, the NIR laser (Ti:Sapphire, operating at 80MHz with *∼* 140 fs pulses, see description of associated methods below) of this two-photon microscope is scanned at low power to excite two-photon fluorescence along Lissajous curves (Fig. S1a). Lissajous scans are single harmonic trajectories that do not generate high harmonic components like a raster scan. They are ideal to make fast and fractional scans of the focal plane. Only 8% of voxels are scanned in the Lissajous trajectories, leading to a more than order of magnitude speed-up with respect to a full, raster scan. We did not investigate systematically the capacity of the surface estimation to handle even sparser Lissajous (reducing *m* and *n*, in equation 5 and 6). It was demonstrated previously that it has the potential to work down to 0.1% of scanned territories^28^. Figure S1b shows the signal obtained form such a Lissajous scan. The structure of interest is hardly recognizable with the human eye. Figs S2a-c shows respectively: a-the detected bright voxels in the sampled volume that are candidate for being GFP lying on the surface of interest; b-the classification of inliers and outliers obtained with the proposed approach; c-the surface obtained after interpolation of the inliers. We observed that the estimated surface matches very well with the epithelium. For example, after performing a ground truth full-scan of sample space, and keeping only the signal from a thin shell of thickness 3*µm* around the estimated surface, a normal cad:GFP image is obtained (Fig. S1c). Thus, the shell could be properly estimated from the fast Lissajous scans, despite the fact that the non-linear process and the low numerical aperture used on this particular setup (NA=0.75) led to a degradation of the signal to noise ratio and resolution compared to the signal generated by the spinning disc confocal. While the peak pulse power involved in this particular set-up did not allow for plasma-induced severing, the demonstrated surface estimation should allow for an efficient implementation of adaptive 3D-scans on stand alone NIR set-up with higher peak-power.

### Methods for Single-point scanning two photon imaging with Lissajous trajectories

In order to test our surface estimation with a single-point scanning two photon microscope, we used a femtosecond (*∼* 140 fs) Ti:Sapphire pulsed laser running at 80 MHz (Chameleon, Coherent) tuned to *∼* 930 nm to excite the two-photon fluorescence of GFP. The laser illuminates a pair of galvanometer scanning mirrors (6215HM60, Cambridge Technology). In turn, scanning mirrors are conjugated to the back focal plane of an objective lens (CFI Plan Apochromat Lambda 20*X* NA=0.75, Nikon) mounted on a commercial microscope stand (Eclipse Ti-U, Nikon) with piezo drive (PIFOC, Physik Instrumente) for the precise Z position control. The same objective lens is used to collect the fluorescence signal generated in a sample (*epi* geometry) and this signal is registered by a photo-multiplier tube (R9110, Hamamatsu). A set of dielectric spectral filters are used to separate the fluorescence radiation at the spectral range of 510 *−* 540 nm from the excitation wavelength: a dichroic mirror T770SPXR (AHFanalysentechnik, AG), a shortpass filter (ET750sp-2p8, Chroma Technology.) and a bandpass filter (510AF23 XF3080, Omega Optical incorporated). Signal acquisition and galvanometer mirrors are synchronized and controlled by an acquisition card (DAQ USB-6351, National Instruments) with use of a custom developed MATLAB code. Change of the deflection angle of the galvanometer mirrors causes the focused beam translation in the sample. The focus trajectories are based on Lissajous curves and defined the following way:

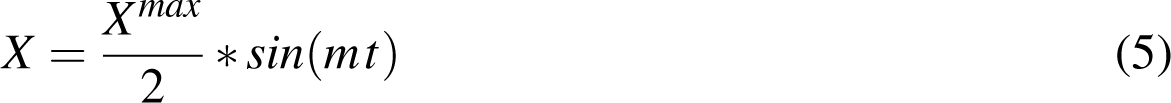

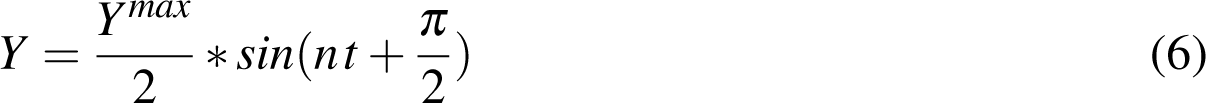

where *X^max^*and *Y^max^* is the field of view size in the *x* and *y* direction, respectively, *t* is a parameter that varies from 0 to 2*π*, *n* and *m* are integer parameters that define the sparsity of the scans. Several pairs of of *n* and *m* are used during acquisition depending on how dense the required scan is: [*n* = 41*, m* = 43], [*n* = 71*, m* = 73], [*n* = 101*, m* = 103]. To estimate the surface with the Lissajous scans, the same procedure is applied than with the 3D stacks from the spinning disc (see methods). Because the Lissajous trajectory is already a a fractional sampling of space, we take for Ω_0_ the full Lissajous trajectory. The intensity distribution from the two-photon Lissajous scans is quite distinct from a Gaussian curve or heavy tailed Gaussian. Although further studies on the noise characteristic could be performed, we found empirically that *T ≈* 8 worked for most instances.

**FIG. S1.**
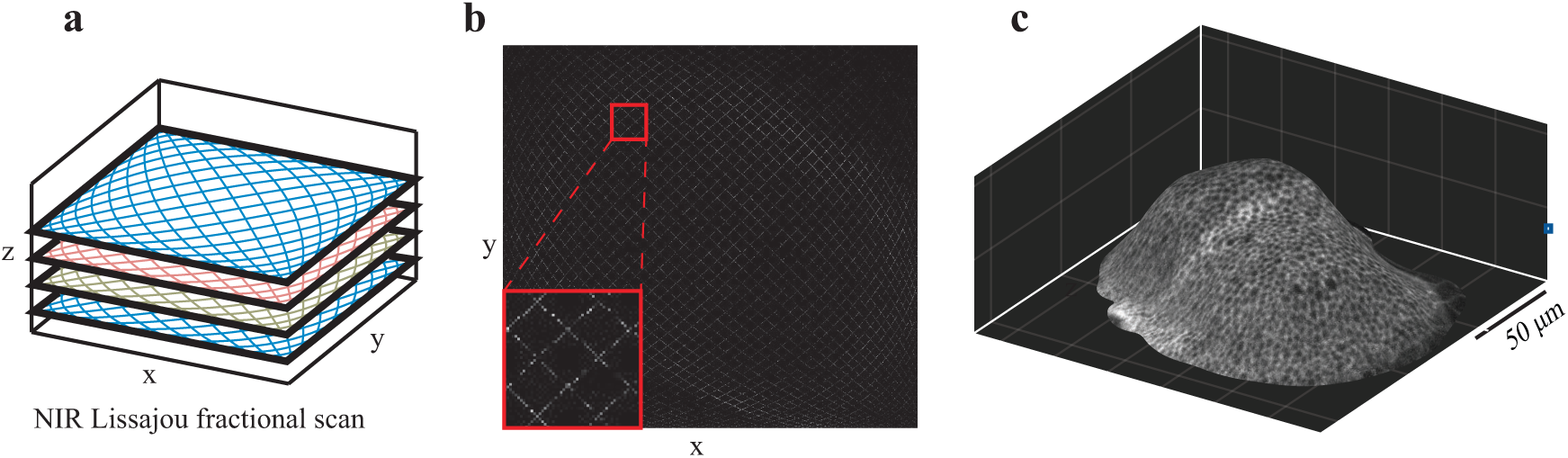
Rapid surface estimation in single-point scanning. To rapidly estimate the surface of interest using only a NIR laser, we perform Lissajous scans, which explores a limited number of voxels within the sample space (a: scheme of the scanned path. Lissajous trajectories following eqs. 5,6 are repeated at different planes; b: collected fluorescence on one plane). (c) In order to verify the quality of the estimated surface with Lissajous scans, we perform a ground truth full-scan of sample space and we only display the voxels located inside a thin shell of thickness 3*µm* around the estimated surface. The fluorescent cad:GFP signal is well encapsulated by the shell.

**FIG. S2.**
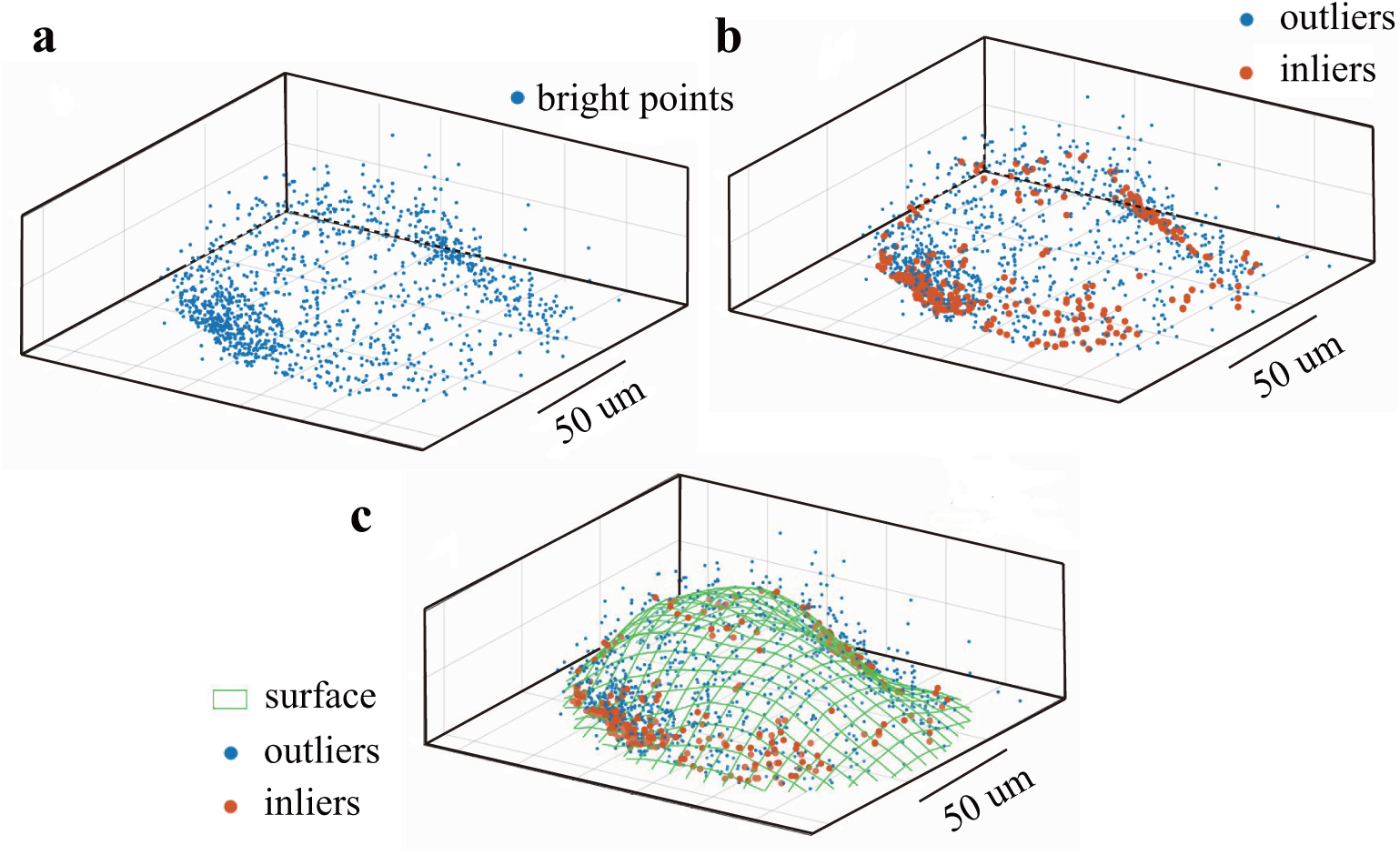
Steps of surface estimation. All data are taken from the surface estimation with Lissajous scans from Figure S1c-f. (a) Bright points detection on the Lissajous prescan after correction for inhomogeneities (eq. 1). (b) classification of bright points into inliers and outliers using the proposed approach. (c) surface estimation form the interpolation of inliers using spline fitting.

**FIG. S3.**
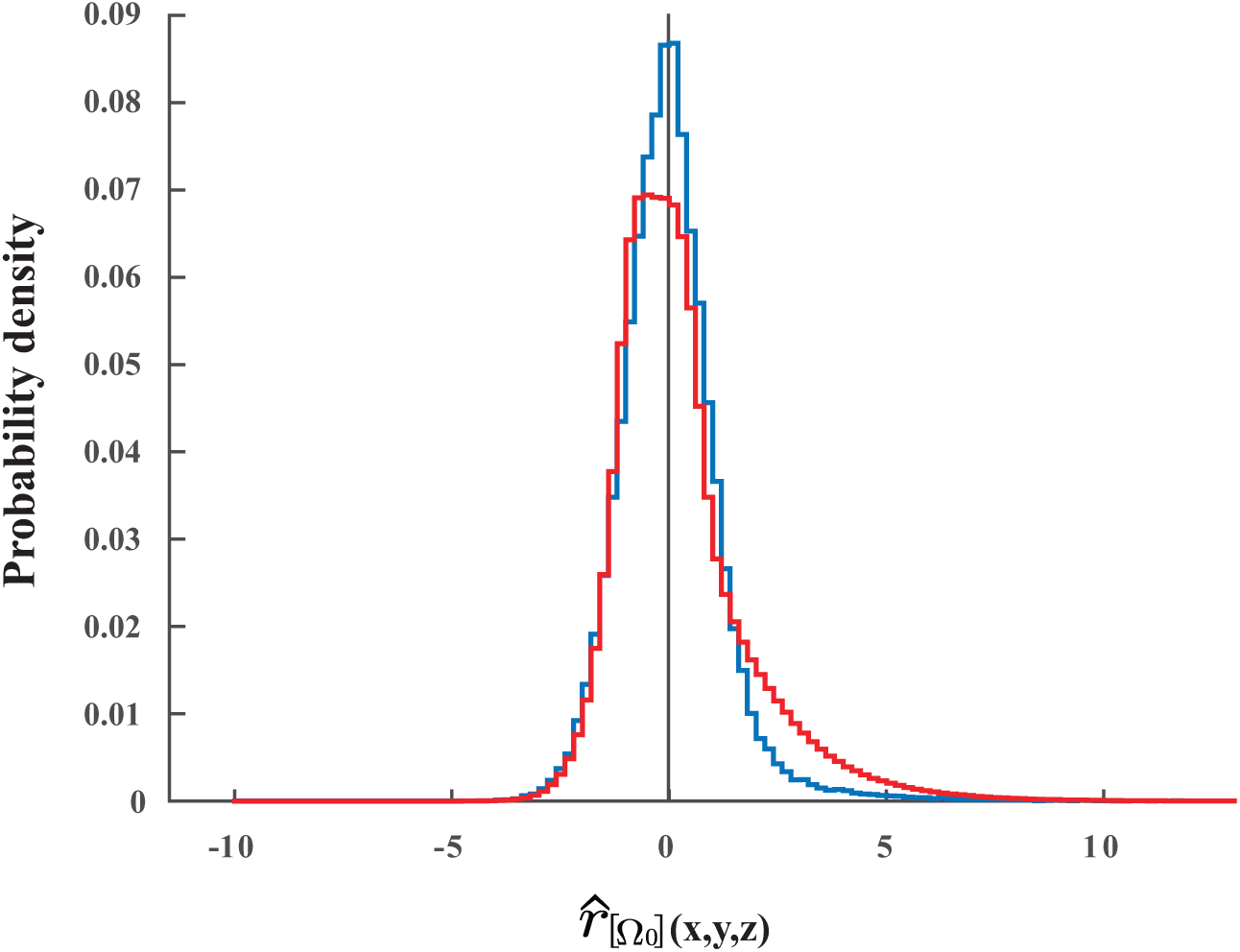
Intensity distributions after correction of inhomogeneities. Distribution of the normalized signal introduced by equation 1 in linear fluorescence stacks and in the two photon fluorescence case. In the spinning disc case (in blue), the distribution is symmetric and gaussian distributed with a small heavy tail at high intensities. In the two photon case (red), the distribution is much less symmetric.

**FIG. S4.**
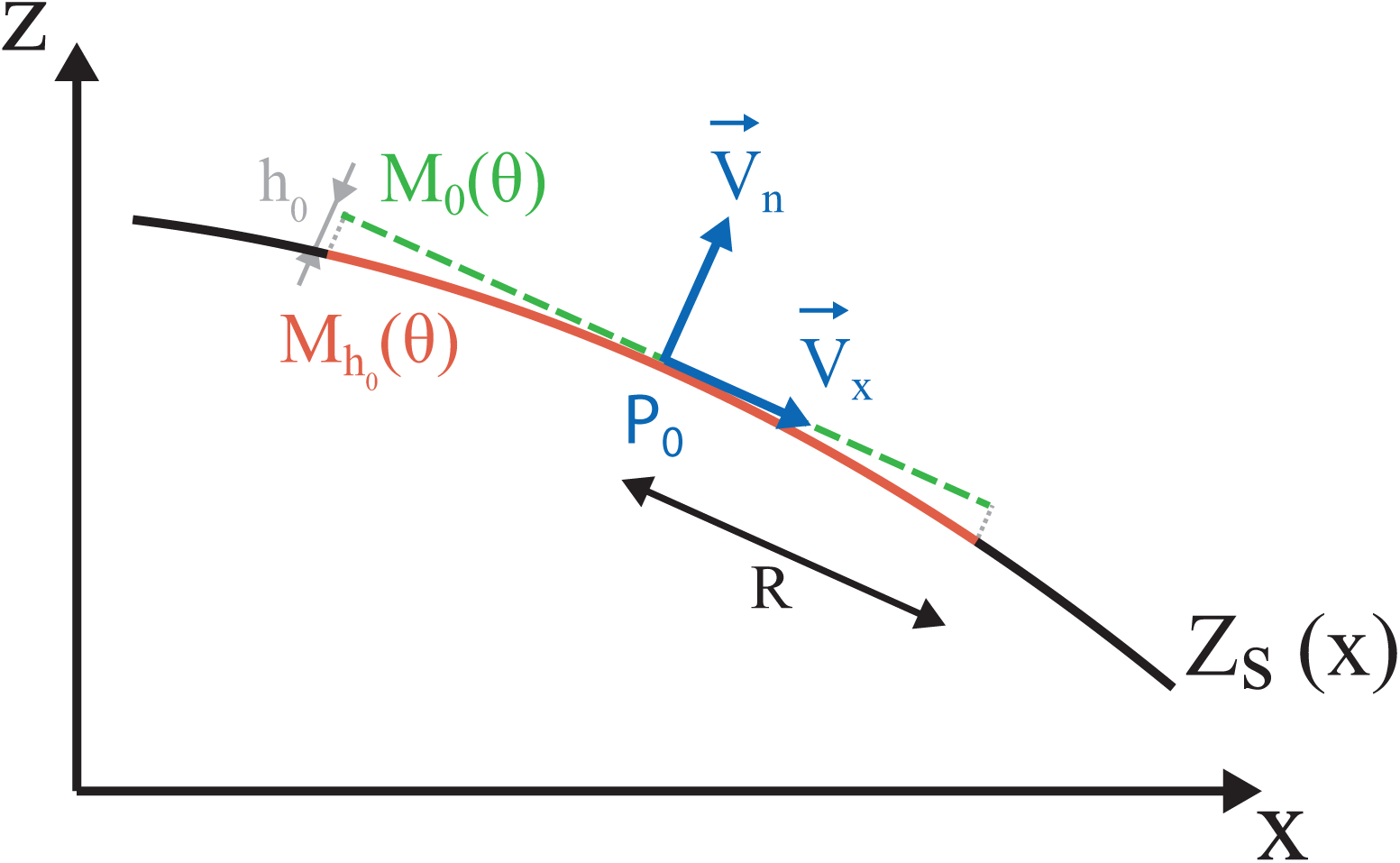
Geometry of dissection on a curved surface. (a) Quadratic surface with local tangent and normal in blue. The green dotted line outlines a strictly circular trajectory, parallel to the local tangent. We project this ring on the surface *Z_s_* (red curve) for an optimal dissection process. The projection is performed along the normal of the surface (*V_n_*). One point of the circular trajectory (*M_θ_* (*θ*)) and its projection (*M_hθ_* (*θ*)), discussed in the method, are also shown.

